# Characterization of deep neural network features by decodability from human brain activity

**DOI:** 10.1101/424168

**Authors:** Tomoyasu Horikawa, Shuntaro C. Aoki, Mitsuaki Tsukamoto, Yukiyasu Kamitani

**Affiliations:** ATR Computational Neuroscience Laboratories, 2-2-2 Hikaridai, Seika, Soraku, Kyoto 619-0288, Japan; Graduate School of Informatics, Kyoto University, Yoshida-honmachi, Sakyo-ku, Kyoto 606-8501, Japan

## Abstract

Achievements of near human-level performances in object recognition by deep neural networks (DNNs) have triggered a flood of comparative studies between the brain and DNNs. Using a DNN as a proxy for hierarchical visual representations, our recent study found that human brain activity patterns measured by functional magnetic resonance imaging (fMRI) can be decoded (translated) into DNN feature values given the same inputs. However, not all DNN features are equally decoded, indicating a gap between the DNN and human vision. Here, we present a dataset derived through the DNN feature decoding analyses including fMRI signals of five human subjects during image viewing, decoded feature values of DNNs (AlexNet and VGG19), and decoding accuracies of individual DNN features with their rankings. The decoding accuracies of individual features were highly correlated between subjects, suggesting the systematic differences between the brain and DNNs. We hope the present dataset will contribute to reveal the gap between the brain and DNNs and provide an opportunity to make use of the decoded features for further applications.

## Background & Summary

Building models that achieve human-level performance has motivated researchers to construct computational models that mimic the architectural and representational properties of the human brain. Adopting the hierarchical architecture of the human visual system, deep neural networks (DNNs) have demonstrated utility in various applications, including object recognition in computer vision, where near human-level performances are achieved. This achievement has led to many comparative studies on the similarity between the brain and DNNs, providing empirical support for the correspondence between the hierarchical representations of the brain and DNNs^1-8^.

On the basis of the hierarchical representational similarity between the brain and DNNs, our recent study demonstrated that human brain activity measured by functional magnetic resonance imaging (fMRI) can be decoded (translated) into DNN feature values^6^. Combining those decoded DNN features and techniques developed with DNNs, recent work has started to develop new technologies to read out richer contents in the brain as demonstrated in the generic decoding of seen, imagined, and dreamed objects^6,7^ and in the reconstruction of seen and imagined images^9^. As exemplified by these studies, decoding of DNN features from brain activity patterns can then provide opportunities to develop new technologies for further applications in brain machine interfacing.

In addition to the capability of the DNN feature decoding approach as a generative model of DNN signal patterns from the brain, the decoding approach also has an advantage allowing to characterize individual DNN units in terms of their decodability from brain activity patterns. Our decoding analysis of DNN features showed that not all DNN feature units were equally decoded^6^, indicating a gap between the DNN and human vision. Thus, evaluating the decodability of individual DNN units will help to further elucidate finer levels of representational similarity between the brain and DNNs, enabling to select highly decodable features for further analyses^10^.

In this report, we present a dataset derived through the DNN feature decoding analyses from human brain activity patterns (Figure 1). The dataset consists of fMRI signals measured while subjects viewed natural images, DNN feature values of all individual units decoded from the measured brain activity patterns, and decoding accuracies of individual units with their rankings among units.

**Figure 1.**
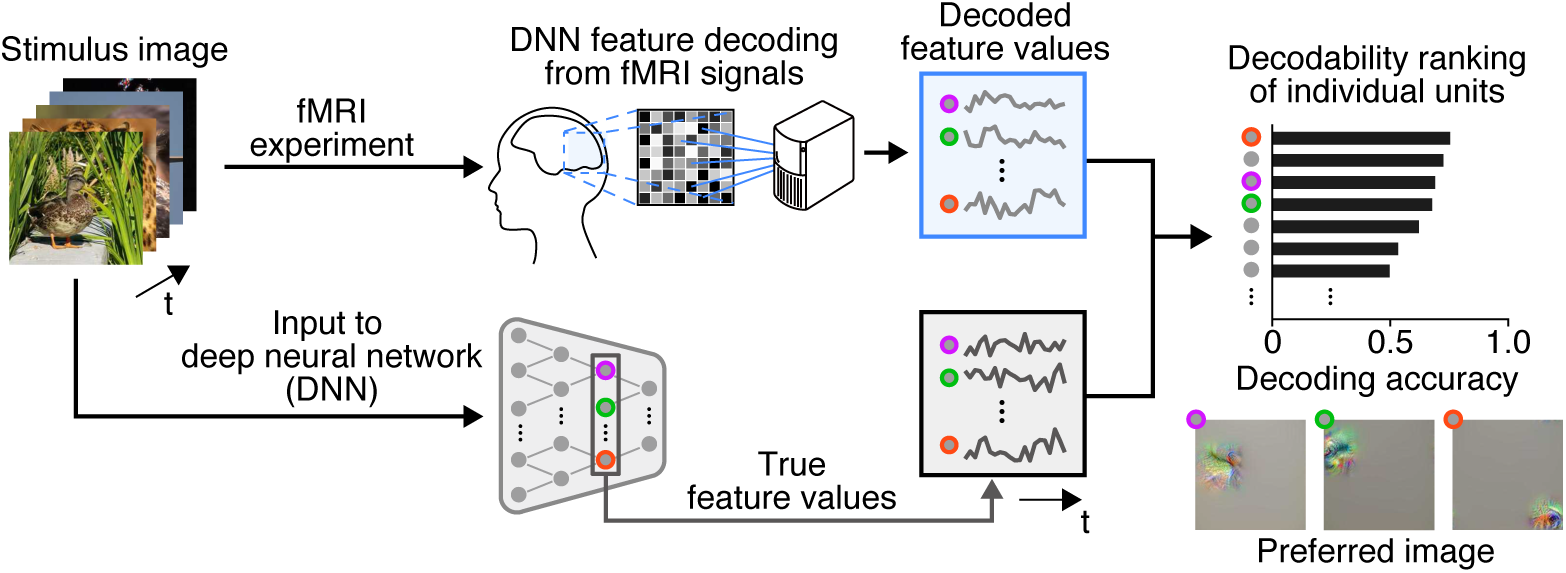
Overview of the data generation procedures. Stimulus images were presented to human subjects in the fMRI experiments to collect fMRI signals. DNN feature decoders were; first tuained to deco de DNN feature values of presented i mages from the training UMRI dat a, and were applied to test fMRI data for producing sequences of decoded feature values for all DNN units. The sam e stimulus images were also provided to DNNs as inputs and sequen ces of DNN feature values were computed for ell DnN units. /or each individual DINN unit, decoding accuracy (or “decodability”) was evaluated using Pearson correlation coefficient be twee n the sequences of decod ed and tr ue featude nalues. The eatimated d ecodability wa s used to rank DNN units within each DNN layer. Examples of preferred image of high rank units are shown at the right bottom.

The fMRI dataset was originally collected in Horikawa and Kamitani (2017a)^6^, which consisted of fMRI signals from five subjects measured while the subjects viewed sequences of natural images (image presentation experiment). This image presentation experiment had two sessions: a training image session and a test image session. Data from the training and test image sessions consisted of fMRI responses to a total of 1,200 and 50 images (“training” and “test” datasets).

The fMRI dataset was used to generate decoded DNN features for individual subjects. Using two types of DNN models, AlexNet^11^ and VGG19^12^, we first computed DNN feature values from the images presented in the fMRI experiments. We then trained a set of statistical linear regression models (decoders) to predict DNN feature values of presented images using visual cortical activity patterns of the training dataset. The trained decoders were applied to the test dataset to produce decoded feature values of the DNNs for the 50 test images for all individual DNN units.

The “decodability” of the individual DNN units were then evaluated for individual subjects. For each DNN unit, a Pearson correlation coefficient was calculated between a sequence of decoded feature values and that of true feature values for the presented 50 test images. Then, the rankings of the decodability were estimated among a set of units within each DNN layer.

Our validation analysis showed that while the decodability was largely varied across units, they were highly correlated across subjects for most DNN layers of the tested DNN models. The results indicate systematic differences between the DNN and the human brain in representing visual images.

To summarize, the present dataset contains a set of resources that is made use of for the DNN feature decoding and for further analyses. We hope that this dataset will offer opportunities to the neuroscience and computer science communities for developing new brain-DNN hybrid applications based on decoded features and for comparative studies aiming at revealing the gap between the brain and DNNs.

## Methods

The data used in this study comes from a previous study performed at our laboratory^6^. According to the journal policy, here, we provide a self-contained description of the subjects, datasets, and preprocessing of the MRI data for the main experiments to make it possible to understand and reproduce the experiments and analyses without referring to associated publications.

### Subjects

Five healthy subjects (one female and four males, aged between 23 and 38) participated in this study. All subjects had normal or corrected-to-normal visual acuity, and had substantial experience participating in fMRI experiments. All studies were performed with the written informed consent of the subjects, and were approved by the Ethics Committee of ATR.

### Stimuli

The stimuli consisted of sequences of natural images collected from an online image database ImageNet^13^ (2011, fall release). We first selected 200 representative object categories (synsets) from the database, and then randomly assigned them to 150 training and 50 test categories. Eight or one images were selected from those training and test categories, respectively. The selected images were cropped to the center.

### Experimental design

The subjects participated in an image presentation experiment, a retinotopy experiment and a functional localizer experiment. All visual stimuli were rear-projected onto a screen in an fMRI scanner bore using a luminance-calibrated LCD projector. Data from each subject were collected over multiple scanning sessions spanning approximately 2 months for the image presentation experiment. On each experiment day, one consecutive session was conducted for at most 2 hours. Subjects were given adequate time for rest between runs (every 3^~^10 minutes), and were allowed to take a break or stop the experiment at any time.

The image presentation experiment consisted of two distinct types of sessions: training image sessions and test image sessions, each of which consisted of 24 and 35 separate runs (9 minutes 54 seconds for each run), respectively. Each run contained 55 stimulus blocks consisting of 50 blocks with different images and 5 randomly interspersed repetition blocks where the same image as in the previous block was presented. In each stimulus block an image (12 × 12 degrees of visual angle) was flashed at 1 Hz for 9 seconds. Images were presented at the center of the display with a central fixation spot. The color of the fixation spot changed from white to red for 0.5 seconds before each stimulus block began to indicate the onset of the block. Extra 33-second and 6-second rest periods were added to the beginning and end of each run, respectively. Subjects were instructed to maintain steady fixation on the fixation spot throughout each run, and performed a one-back repetition detection task on the images, responding with a button press for each repetition to maintain their attention on the presented images (mean task performance across five subjects; sensitivity = 0.930; specificity = 0.995). In the training image session, a total of 1,200 images from 150 categories (eight images from each category) were each presented once. In the test image session, a total of 50 images from 50 object categories (one image from each category) were presented 35 times each. The presentation order of the categories was randomized across runs.

The retinotopy experiment was performed by following the conventional protocol^14,15^ using a rotating wedge and an expanding ring of a flickering checkerboard. The data were used to delineate the borders between each visual cortical area, and to identify the retinotopic map (V1–V4) on the flattened cortical surfaces of individual subjects.

The functional localizer experiment was performed to identify the lateral occipital complex (LOC)^16^, fusiform face area (FFA)^17^, and parahippocampal place area (PPA)^18^ for each individual subject. The localizer experiment consisted of four to eight runs (varied across subjects) and each run contained 16 stimulus blocks. In this experiment, intact or scrambled images (12 × 12 degrees of visual angle) from face, object, house, and scene categories were presented at the center of the screen. Each of eight stimulus types (four categories × two conditions) was presented twice per run. Each stimulus block consisted of a 15-second intact or scrambled image presentation. The intact and scrambled stimulus blocks were presented successively (the order of the intact and scrambled stimulus blocks was randomized), followed by a 15-second rest period consisting of a uniform gray background. Extra 33- second and 6-second rest periods were added to the beginning and end of each run, respectively. In each stimulus block, 20 different images of the same type were presented for 0.3 seconds, followed by an intervening blank screen of 0.45 seconds.

### MRI acquisition

MRI data were collected using 3.0-Tesla Siemens MAGNETOM Trio A Tim scanner located at the ATR Brain Activity Imaging Center. An interleaved T2*-weighted gradient-EPI scan was performed to acquire functional images to covering the entire brain (image presentation and localizer experiments: TR, 3,000 ms; TE, 30 ms; flip angle, 80 deg; FOV, 192 × 192 mm; voxel size, 3 × 3 × 3 mm; slice gap, 0 mm; number of slices, 50) or the entire occipital lobe (retinotopy experiment: TR, 2,000 ms; TE, 30 ms; flip angle, 80 deg; FOV, 192 × 192 mm; voxel size, 3 × 3 × 3 mm; slice gap, 0 mm; number of slices, 30). T2-weighted turbo spin echo images were scanned to acquire high-resolution anatomical images of the same slices used for the EPI (image presentation and localizer experiments: TR, 7,020 ms; TE, 69 ms; flip angle, 160 deg; FOV, 192 × 192 mm; voxel size, 0.75 × 0.75 × 3.0 mm; retinotopy experiment: TR, 6,000 ms; TE, 57 ms; flip angle, 160 deg; FOV, 192 × 192 mm; voxel size, 0.75 × 0.75 × 3.0 mm). T1-weighted magnetization-prepared rapid acquisition gradient-echo (MP-RAGE) fine-structural images of the entire head were also acquired (TR, 2,250 ms; TE, 3.06 ms; TI, 900 ms; flip angle, 9 deg, FOV, 256 × 256 mm; voxel size, 1.0 × 1.0 × 1.0 mm).

### MRI data preprocessing

The first 9-second scans for experiments with TR = 3 seconds (three volumes; image presentation and localizer experiments) and 8-second scans for experiments with TR = 2 seconds (four volumes; retinotopy experiment) of each run were discarded to avoid MRI scanner instability. The acquired fMRI data underwent three-dimensional motion correction using SPM5 (http://www.fil.ion.ucl.ac.uk/spm). The data were then coregistered to the within-session high-resolution anatomical image of the same slices used for EPI and subsequently to the whole-head high-resolution anatomical image. The coregistered data were then reinterpolated by 3 × 3 × 3 mm voxels.

For the data from the image presentation experiment, data samples were created by first regressing out nuisance parameters from each voxel amplitude for each run, including a constant baseline, a linear trend, and temporal components proportional to the six motion parameters calculated from the SPM motion correction procedure. The data were then despiked to reduce extreme values (beyond ± 3SD for each run). The voxel amplitudes were then averaged within each 9-second stimulus block (three volumes) after shifting the data by 3 seconds (one volume) to compensate for hemodynamic delays.

### Region of interest (ROI) selection

V1, V2, V3, and V4 were delineated by the standard retinotopy experiment^14,15^. The data from the retinotopy experiment were transformed to Talairach coordinates and the visual cortical borders were delineated on the flattened cortical surfaces using BrainVoyager QX (http://www.brainvoyager.com). The voxel coordinates around the gray-white matter boundary in V1–V4 were identified and transformed back into the original coordinates of the EPI images. The LOC, FFA, and PPA were identified using conventional functional localizers^16-18^. The data from the functional localizer experiment were analyzed using SPM5. The voxels showing significantly higher responses to intact object, face, or scene images than to corresponding scrambled images (two-sided *t*-test, uncorrected *P* < 0.05 or 0.01) were identified, and defined as LOC, FFA, and PPA, respectively. A contiguous region covering the LOC, FFA, and PPA was manually delineated on the flattened cortical surfaces, and the region was defined as the “higher visual cortex” (HVC). Voxels from V1-V4 and the HVC were combined to define the “visual cortex” (VC). In the regression analysis, voxels showing the highest correlation coefficient with the target variable in the training dataset were provided to decoders constructed for individual feature units (with a maximum of 500 voxels).

### Deep neural networks (DNNs)

We used the *Caffe* implementation of the *AlexNef*^11^ and *VGG19*^12^ deep neural network models (available from https://github.com/BVLC/caffe/), both of which were pre-trained with images in ImageNet (Deng et al., 2009) to classify 1,000 object categories. The AlexNet consisted of five convolutional layers (conv1, conv2, conv3, conv4, and conv5) and three fully-connected layers (fc6, fc7, and fc8). The VGG19 model consisted of a total of sixteen convolutional layers (conv1_1, conv1_2, conv2_1, conv2_2, conv3_1, conv3_2, conv3_3, conv3_4, conv4_1, conv4_2, conv4_3, conv4_4, conv5_1, conv5_2, conv5_3, conv5_4), and three fully-connected layers (fc6, fc7, and fc8). The outputs from the units in each of the DNN layers (immediately after convolutional or fully connected layers, before rectification) were used as target variables in the following feature decoding analysis.

### Deep neural network feature decoding

We used a set of linear regression models to construct multivoxel decoders to decode DNN feature values of a seen image from a fMRI activity pattern. In this study, we used the sparse linear regression (SLR) algorithm^19^ that can automatically select the important voxels for prediction. In our analysis, a single regression model (decoder) was constructed to predict feature values of a single DNN unit. In the following, we explain the regression model for a single DNN unit. We individually trained multiple models for predicting feature values of all DNN units in the tested DNN layers and models.

Given an fMRI data sample **x** = {*x*_1_, …,*x_d_*}^T^ consisting of *d* voxels’ activities as input, the regression function can be expressed by

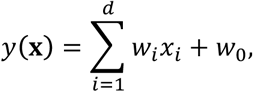

where *x_i_* is the fMRI amplitude of the voxel *i*, *w_i_* is the weight of voxel *i*, and *w*_0_ is the bias. For simplicity, the bias *w*_0_ is included into the weight vector such that **w** = {*w*_0_, …,*w_d_*}^T^. The dummy variable *x*_0_ = 1 is introduced into the data such that **x** = {*x*_0_,…, *x_d_*}^T^. Using this regression function, we modelled the activity of a DNN unit as a target variable *t* that is explained by the regression function *y*(**x**) with additive Gaussian noise as described by

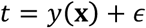

where *ϵ* is a zero mean Gaussian random variable with noise precision *β*.

Given a training data set, SLR computes the weights for the regression function such that the regression function optimizes an objective function. To construct the objective function, we
first express the likelihood function by

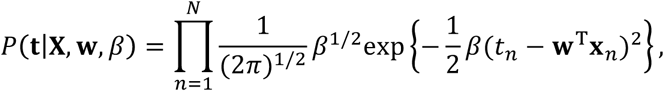

where *N* is the number of samples, **X** is an *N* × (*d* + 1) fMRI data matrix whose *n*th row is the *d* + 1-dimensional vector **x**_*n*_, and **t** = {*t*_1_, …, *t_N_*}^T^ are the samples of a DNN unit.

To introduce sparsity into the weight estimation, we performed Bayesian parameter estimation, and adopted the automatic relevance determination (ARD) prior^19^. We considered the estimation of the weight parameter **w** given the training data sets {**X, t**}. We assumed a Gaussian distribution prior for the weights **w** and non-informative priors for the weight precision parameters **α** = {*α*_0_,… *α_d_*}^T^ and the noise precision parameter *β*, which are described as

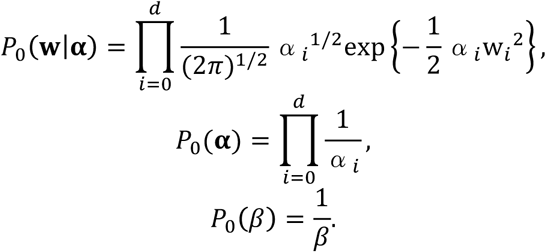

In the Bayesian framework, we considered the joint probability distribution of all the estimated parameters, and the weights can be estimated by evaluating the following joint posterior probability of **w**:

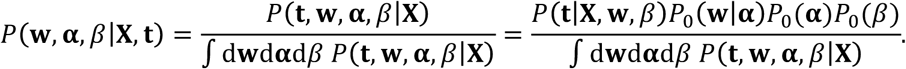

Given that the evaluation of the joint posterior probability *P*(**w**,**α**,*β*|**X**,**t**) is analytically intractable, we approximated it using the variational Bayesian method^19-21^. While the results presented in this manuscript were obtained from the models with the ARD prior, qualitatively similar results were obtained using other regression models (e.g., ordinary least square regression model).

We trained linear regression models that decode feature values of individual feature units for seen images given fMRI samples in the training image session. For test dataset, fMRI samples corresponding to the same images (35 samples for each of the 50 test images) were averaged across trials to increase the signal to noise ratio of the fMRI signals. Using the learned models, we decoded feature values of seen images from averaged fMRI samples. Feature decoding accuracy of each DNN unit was evaluated by the Pearson correlation coefficient between the true and decoded feature values of each feature unit. The estimated correlation coefficients (“decodability”) from individual subjects and their averages were ranked within each DNN layers and models separately. We assigned *nan* values to the decodability and their ranks for units not showing any responses (DNN signals) to images in the training or test datasets.

### Preferred images of individual units

We used the activation maximization technique to generate preferred images of individual units in each DNN layer^22-25^. Generating preferred images starts from a random image and optimizes the image to maximally activate a target DNN unit by iteratively calculating how the image should be changed via backpropagation. This analysis was implemented using custom software written in MATLAB based on Python codes provided in a series of blog posts (Mordvintsev, A., Olah, C., Tyka, M., DeepDream—a code example for visualizing Neural Networks, https://github.com/google/deepdream, 2015; Øygard, A. M., Visualizing GoogLeNet Classes, https://github.com/auduno/deepdraw, 2015).

### Code availability

The code for the DNN feature decoding is available at https://github.com/KamitaniLab/GenericObjectDecoding. Both MATLAB and Python scripts are included in the repository. We also provide a Python API to download and extract data from Figshare (Data Citation2: Figshare https://doi.org/10.6084/m9.figshare.6269321), and jupyter notebooks for usage example of the data at https://github.com/KamitaniLab/brain-decoding-datasets. All code is available without any access restrictions.

## Data Records

### Experimental data

All data produced from the MRI experiments are hosted at OpenNeuro (Data Citation 1: OpenNeuro ds001246). The dataset is based on Brain Imaging Data Structure (BIDS)^26^. All MRI images are saved as NIfTI files.

The data repository contains five directories for the five subjects (*sub*-*01* to *sub*-*05*). Each directory consists of several subdirectories that include MRI data in a single scanning session. *ses*-*anatomy* contains a defaced Tl-weighted anatomical reference image for the individual subject. *ses*-*perceptionTraining*^*^ and *ses*-*perceptionTest*^*^ directories include fMRI images collected in the training and image presentation experiments, respectively (Table 1). fMRI images from a single run are stored in a single 4-D NIfTI file. Each run is accompanied by a task event file, which describes experiment information such as timing of trials, presented stimuli, and subject’s response time (Table 2). The session directories also contain a T2- weighted anatomical image obtained in the session. Binary mask images for ROIs used in the analysis (see above) are placed in *sourcedata*/<*subject*>/*anat* directories (Table 3).

**Table 1:**
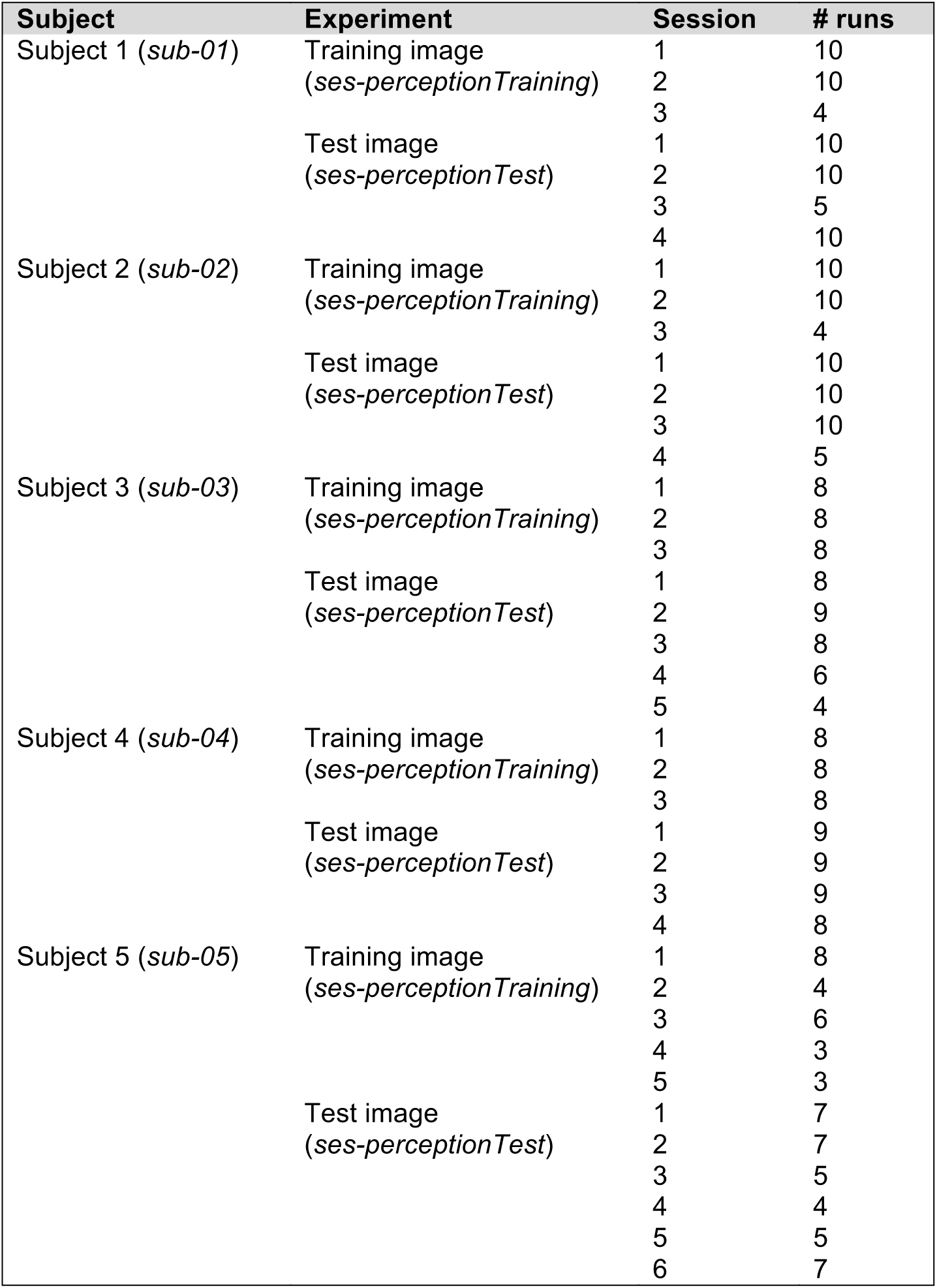
Summary of the experimental data.

**Table 2:**
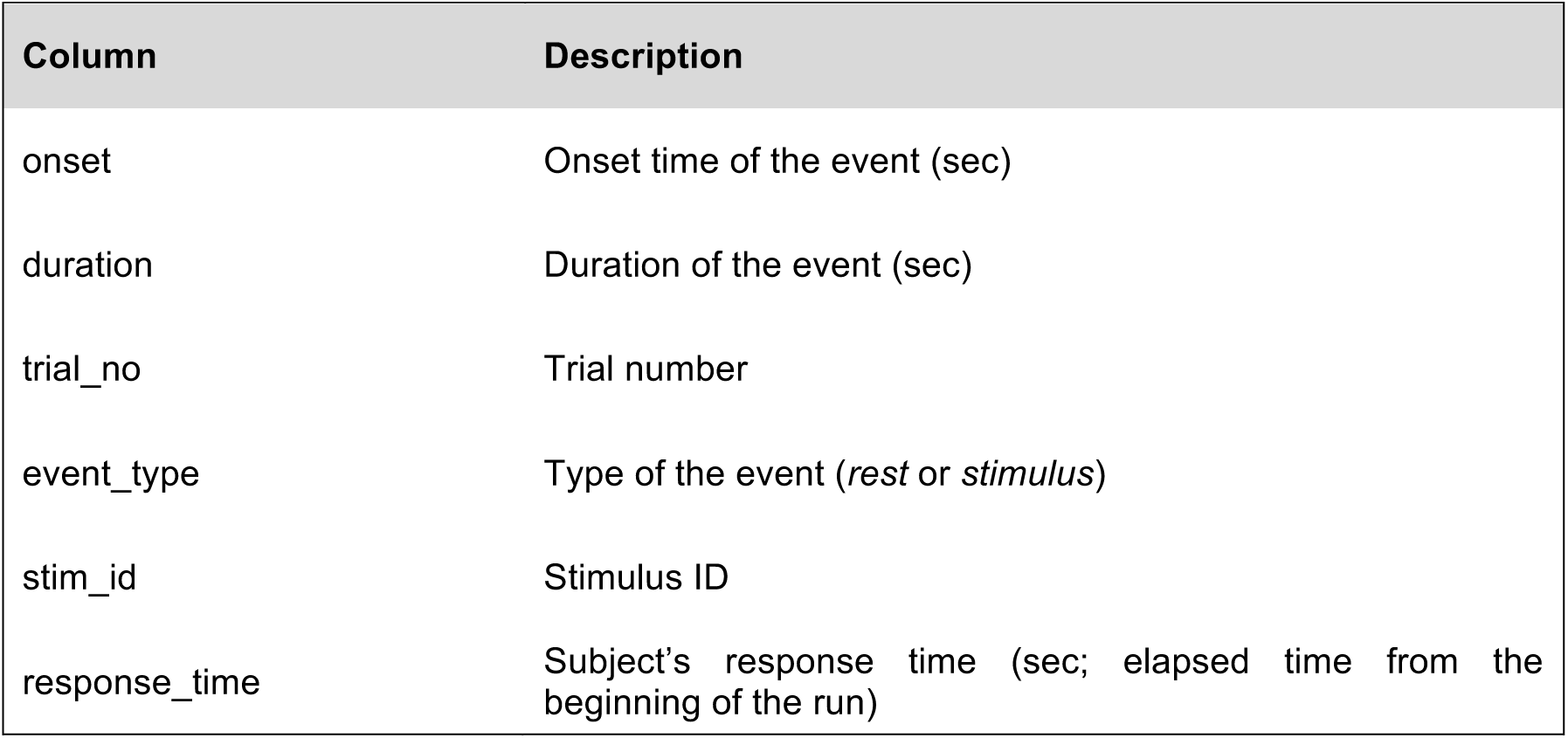
Columns in task event files for the image presentation experiments.

**Table 3:** ROI mask images included in the dataset.

| File name | ROI |
| --- | --- |
| sub-*_mask_LH_V1.nii.gz | Left V1 |
| sub-*_mask_RH_V1.nii.gz | Right V1 |
| sub-*_mask_LH_V2.nii.gz | Left V2 |
| sub-*_mask_RH_V2.nii.gz | Right V2 |
| sub-*_mask_LH_V3.nii.gz | Left V3 |
| sub-*_mask_RH_V3.nii.gz | Right V3 |
| sub-*_mask_LH_hV4.nii.gz | Left V4 |
| sub-*_mask_RH_hV4.nii.gz | Right V4 |
| sub-*_mask_LH_LOC.nii.gz | Left LOC |
| sub-*_mask_RH_LOC.nii.gz | Right LOC |
| sub-*_mask_LH_FFA.nii.gz | Left FFA |
| sub-*_mask_RH_FFA.nii.gz | Right FFA |
| sub-*_mask_LH_PPA.nii.gz | Left PPA |
| sub-*_mask_RH_PPA.nii.gz | Right PPA |
| sub-*_mask_LH_HVC.nii.gz | Left higher visual cortex (HVC) |
| sub-*_mask_RH_HVC.nii.gz | Right higher visual cortex (HVC) |

In the task event files, stimuli are represented by a float number, *stimulus_id*, in which the integer part indicates WordNet^27^ ID for the synset (category) and the decimal part indicates image ID. For example, *1518878.005958* represents image *5958* in synset *n01518878* (‘ostrich’). Due to license issues, we do not include the stimulus images in the data repository. A script downloading the stimulus images are available at https://github.com/KamitaniLab/GenericObjectDecoding. Downloaded image files are named as *XXXX*-*YYYY.JPEG*, where *XXXX* and *YYYY* represents the WordNet ID and image ID, respectively (e.g., *n01518878_5958.JPEG*).

### DNN feature and decodability

Decoded feature, true feature, accuracy, and ranking by accuracy are available from Figshare (Data Citation2: Figshare https://doi.org/10.6084/m9.figshare.6269321). All data files are saved as MATLAB (^*^.mat) files and zipped by the DNN and the layer. Naming rule of the ^*^.mat files and size (shape) of the data in the file are summarized in Table4.

**Table 4:**
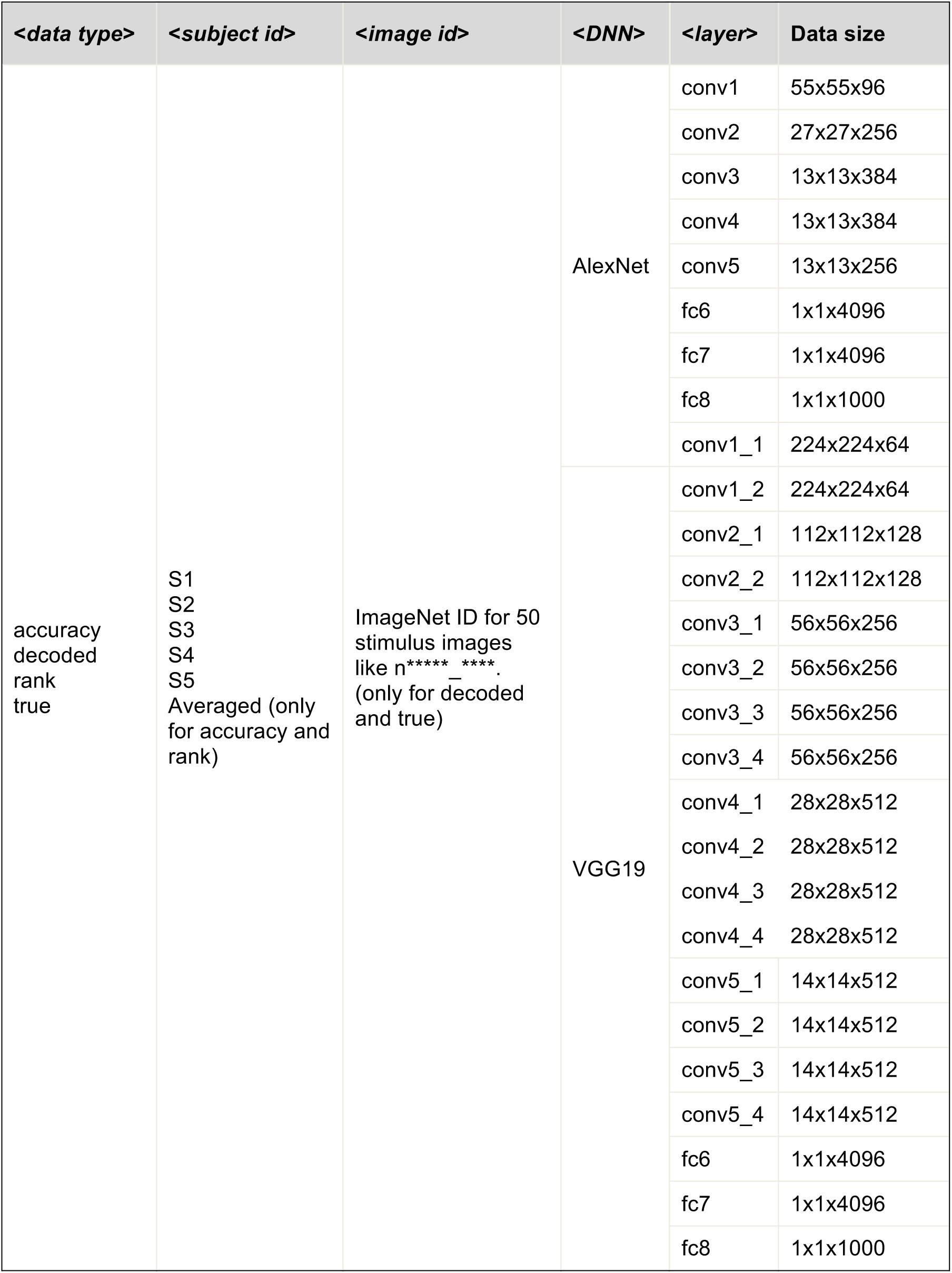
Summary of the DNN feature and decodability datasets. The data files in Figshare are named as “<*data_type*>-<*DNN*>-<*layer*>-<*subject id*>-<*image* id>.mat”. List of each component and size (shape) of the data are shown.

### Decoded DNN feature

The decoded features are saved to a file named like ‘decoded-<*net*>- <*layer*>-<*subject_id*>-<*image_id*>.mat’, where <*net*> takes either “AlexNet” or “VGG19”, <*layer*> takes the layer name of the DNN, <*subject_id*> is the subject ID, and <*image_id*> is the ImageNet ID of the stimulus image. In the ^*^.mat file, an array whose shape is the same as the shape of the output of the <*layer*> layer in the <*net*> DNN model (See Table 4) is saved. The ^*^.mat files are zipped for each DNN and layer to ‘decodedDNN-decoded-<*net*>- <*layer*>.zip’ file and uploaded to Figshare (Data Citation2: Figshare https://doi.org/10.6084/m9.figshare.6269321).

### True DNN feature

The true features are saved for each DNN model, layer, and image, and named as ‘true-<*net*>-<*layer*>-<*image_id*>.mat’. Zipped files for each <*net*> and <*layer*> are uploaded to Figshare.

### Decoding accuracy (“decodability”)

The decoding accuracy for each DNN, layer, and subject are saved in the file named like ‘accuracy-<*net*>-<*layer*>-<*subject_id*>.mat’. In addition, we also created averaged accuracy by subject, ‘accuracy-<*net*>-<*layer*> -Averaged.mat’. Zipped files for each <*net*> and <*layer*> are uploaded to Figshare.

### Decodability ranking

The ranking of feature units by accuracy is provided for each DNN, layer, and subject in a ^*^.mat file named as ‘rank-<*net*>-<*layer*>-<*subject_id*>.mat’. In addition, we also created a file containing average ranking by subject, ‘rank-<*net*>-<*layer*>- Averaged.mat’. Zipped files for each <*net*> and <*layer*> are uploaded to Figshare.

We provide a Python API to download and extract data from Figshare (https://github.com/KamitaniLab/brain-decoding-datasets, see Usage Notes section).

## Technical Validation

In order to validate the quality of the dataset, we first performed feature decoding analysis to decode DNN feature values from fMRI activity patterns, and evaluated the decoding accuracy (“decodability”) of individual DNN units for each subject^6^. After that, we evaluated the consistency of the decodability between multiple subjects to demonstrate the replicability of the results across subjects.

In the feature decoding analysis, decoders were trained to decode DNN unit activities to input image sequences from visual cortical activities measured while the subjects viewed the same sequences of stimulus images using the training dataset (1200 samples). The decoders were individually trained for each unit in the convolutional (5 and 16 layers for AlexNet and VGG19, respectively) and fully-connected layers (3 layers for both of AlexNet and VGG19) of the DNN models (see Table4 for the number of units in each layer). The trained decoders were then applied to an independent test dataset (50 samples) to evaluate the decodability of individual DNN units. The decodability of each DNN unit was evaluated by calculating a correlation coefficient (Pearson correlation) between a pair of feature value sequences from the DNN (true features) and the brain activity of individual subjects (decoded features).

The obtained decodabilities of individual units were further examined for each DNN layer and model, and were compared across subjects. Figure 2a shows the distributions of feature decoding accuracies evaluated for each individual layer of each DNN model (AlexNet and VGG19), in which the decoding accuracy largely varied across units, layers, and models. To assess the degree of consistency of decodability across subjects, we evaluated the unit-by-unit similarity of the decodability between multiple subjects. Figure 2b shows example scatter plots of feature decoding accuracies between two subjects. The decodability of individual units from the two subjects densely distributed along the diagonal axis for most layers, showing positive correlations between the two subjects. Figure 2c shows the mean correlation coefficients across all pair combinations of the five subjects. The decodability between subjects show positive correlation coefficients for all layers of the each of the two DNN models. The results suggest that the feature decoding from the brain can produce replicable results and that the decodability was highly consistent across subjects even at the unit level.

**Figure 2.**
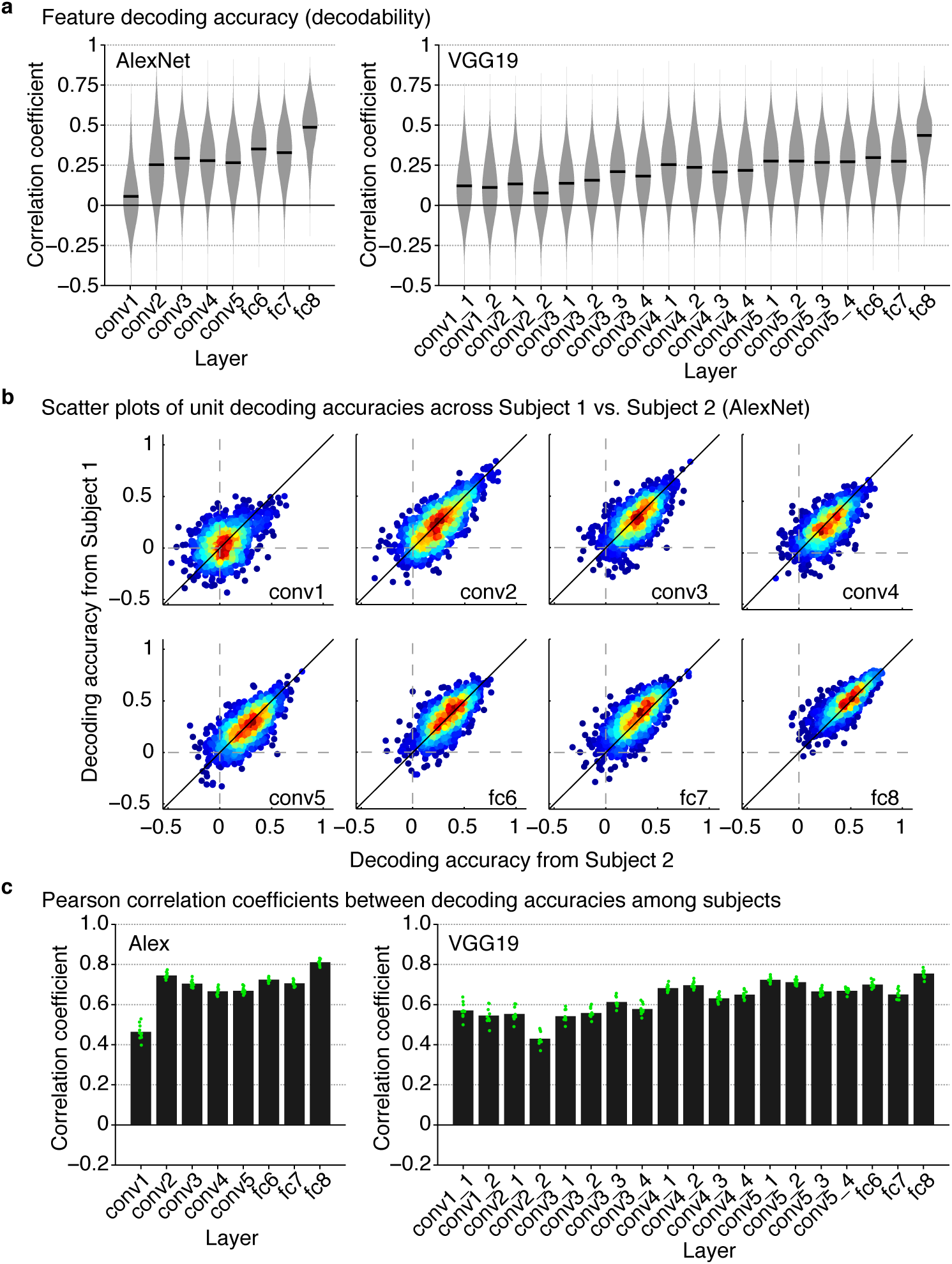
Evaluations of DNN feature decoding. (**a**) Violin plot of feature decoding accuracy for each DNN layer and model. Distributions of the decoding accuracy of all individual units in each DNN layer are shown (pooled across five subjects). Black bars denote mean decoding accuracies averaged across all units and subjects. (**b**) Scatter plots of decoding accuracies of individual DNN units from two subjects (AlexNet). Each dot denotes the decoding accuracy of each DNN unit estimated from Subject 1 (vertical axis) and Subject 2 (horizontal axis). The color of each dot indicates the density of the plotted dots. For visualization purpose, randomly selected subsets of units are shown with a maximum of 1000 units. (**c**) Mean correlation coefficients between decoding accuracies of DNN units from different subjects. Peanson correlation coefficients between decoding accuracies of individual DNN units obtained from different subjects were calculated for all pairs of subjects (10 pairs from 5 subjects). Each dot denotes the correlation coefficients for each pair of subjects.

Taken together, our analyses support the quality of the present dataset as the data showed replicable and consistent results from multiple subjects. The fMRI data made it possible to decode DNN feature values from the brain activity patterns, and the estimated decodability was highly consistent across subjects. Thus, the present dataset could provide an opportunity to utilized for various purposes, including the feature selection in neural encoding and decoding analyses^4,8,10^ and further applications by combining the decoded features with deep neural network technology^6,9^.

## Usage Notes

The experimental data can be downloaded from OpenNeuro (Data Citation 1: OpenNeuro ds001246). To perform DNN feature decoding, the fMRI data need to be preprocessed as described in the Methods section. The head motion correction, functional-anatomical registration in individual anatomical space, and resampling can be conducted with SPM, and the following preprocessing, including regressing-out of nuisance parameters, reduction of extreme values, shifting data, and within-block averaging can be conducted with Brain Decoder Toolbox 2 (https://github.com/KamitaniLab/BrainDecoderToolbox2). The DNN feature decoding analysis can be performed with scripts available at https://github.com/KamitaniLab/GenericObjectDecoding (*analysis_FeaturePredicion.m* for MATLAB and *analysis_FeaturePrediciton.py* for Python). The scripts train feature decoding models with fMRI data in the training image presentation experiments, and predict DNN features from fMRI data in the test image presentation experiments. To feed data to the scripts, the fMRI data must be saved in Brain Decoder Toolbox 2 format.

The decoded DNN features are available at Figshare (Data Citation 2: Figshare https://doi.org/10.6084/m9.figshare.6269321). We provide a Python API for downloading and extracting data from Figshare (https://github.com/KamitaniLab/brain-decoding-datasets). The repository also includes a jupyter notebook that replicates results in Fig. 2.

## Acknowledgements

This research was supported by grants from the New Energy and Industrial Technology Development Organization (NEDO), JSPS KAKENHI Grant number JP26119536, JP15H05920, JP15H05710, JP17K12771 and ImPACT Program of Council for Science, Technology and Innovation (Cabinet Office, Government of Japan).

## Author contributions

T.H. and Y.K. designed the study. T.H. performed experiments and analyses. S.C.A and M.T. arranged and curated datasets. S.C.A. developed utilities for downloading and loading datasets. T.H., S.C.A., M.T. and Y.K. wrote the paper.

## Competing interests

The authors declare no competing interests.

